# Evolutionary Variation of Poxvirus Genome Architecture

**DOI:** 10.1101/2025.10.08.681269

**Authors:** Junior Ayuk Enow, Arkaidy Garber, Betram Jacobs, Grant McFadden, Masmudur M. Rahman

**Affiliations:** Biodesign Center for Personalized Diagnostics, Biodesign Institute, Arizona State University, Tempe, Arizona, 85287, U.S.A.; Biodesign Center for the Mechanism of Evolution, Biodesign Institute, Arizona State University, Tempe, Arizona, 85287, U.S.A.; ASU-Banner Neurodegenerative Disease Research Center, Biodesign Institute, Arizona State University, Tempe, Arizona, 85287, U.S.A.; School of Life Sciences, Arizona State University, Tempe, Arizona, 85287, U.S.A.

## Abstract

Poxviruses (PXV) are large double-stranded DNA viruses (dsDNA) that infect and cause disease in a wide variety of hosts, including humans.. They are studied as models to understand host-specific disease and they have a wide variety of applications in biotechnology, including vaccines, oncolytic virotherapy, and immunotherapy. Despite a nearly fourfold variation in genome sizes across the Poxviridae family, the basic genome architectural features driving their evolution remain poorly understood.

In this study of the summated poxvirus genome sequence database, we report that while only a weak negative correlation exist between viral coding densities and genome sizes, we detected a strong positive correlation between the number of assigned open reading frames (ORF) and genome sizes. We observe an inflection point at a genome size of 229 kb, above or below with the percentage of genes encoding secreted proteins increasing with genome size, and the percentage of predicted cytoplasmic proteins decreasing with genome size. Additionally, we observed a weak positive correlation between the potential of transcript overlap that would be expected to generate dsRNA and genome size.

Unexpectedly, we observed a significant positive correlation between coding density and GC-content in mpox virus isolates (*R*^2^ = 0.799). Further analysis illustrated a rise in both GC-content and coding density of mpox isolates between 2022 and 2024. Codon usage bias was observed to be clustered by virus families, with tryptophan being the least utilized amino acid across all poxviruses.

Protein distribution analysis revealed a right-skewed distribution of poxviral predicted ORFs, with the 150 - 250 amino acid cohort containing the highest number of viral proteins. Finally, we observe a significant positive correlation between ITR sizes and duplicate gene clusters (*R*^2^ = 0.679).

Together, these architectural findings outline the constraints and adaptive pathways governing poxviral evolution and biology.

**Importance:** This study establishes a family genome design rulebook for poxviruses and illustrates how these rules might be linked with their biology and evolution. We demonstrate that genome growth is accompanied by a modest increase in noncoding DNA and an increase in extracellular proteome investment. We further demonstrate that ITR expansions correlate with duplicate gene clusters, rather than genome size, indicating that ITRs serve as hubs of innovation. Our results provide a standard for generating hypotheses and experiments with poxviruses. This work also introduces matrices for within-family viral comparisons.

## Introduction

Poxviruses are large, enveloped, double-stranded DNA (dsDNA) viruses that replicate entirely within the cytoplasm of infected cells and infect a wide range of host species, including insects and vertebrates. They are the causative agents of a wide variety of human and livestock diseases, including smallpox, mpox, and cowpox. Poxviruses are broadly classified into the vertebrate-infecting Chordopoxviruses and the insect-infecting Entomopoxviruses [12, 7, 9, 18, 11].

They are brick-shaped viruses, measuring 220–450 nm in length, 140–260 nm in width, and 140–260 nm in thickness [12]. They contain a linear dsDNA genome covalently linked on both ends [16]. The genome sizes ranges from 120 to 450 kilobases (kb) [12]. The genome is organized such that the centrally located genes are involved in virion morphogenesis and assembly, while the genes situated towards the periphery are essential for modulating the host immune response [17, 18]. The poxviral genome ends in inverted terminal repeats (ITRs) that contain identically, oppositely oriented DNA sequences[3]. During infection, viral genes are transcribed to the left or right, depending on the strand encoding the open reading frame (ORF) [8].

Despite a nearly fourfold variation in poxvirus genome size, a comparison of the genome architectural variation that governs their evolution is lacking. To date, no comprehensive study has addressed the genetic variation in the genome organization within poxviruses or any virus family. Studies describing the within-family genetic diversity of viruses might serve as the foundation of understanding the paths that are evolutionarily open or closed to any viral lineage. It also offers a foundational framework for understanding the cellular biology of viruses, their commonalities, differences, and ultimate fates.

In this study, we performed a systematic analysis of basic poxviral genome features, including predicted ORFs, ORFs’ overlap potential, predicted ORF protein localization, GC content, amino acid usage bias, ITR size, duplicate genes, and genome size. Our findings reveal approximately a 3.75-fold variation in genome size amongst poxviruses. We observed a weak negative correlation between viral coding density and genome size and a linear increase in ORF number with increasing genome size, albeit with considerable variation around the best-fit line. We also report a weak positive correlation between the potential of transcript overlap predicted to generate dsRNA and genome size, again with considerable variation around the regression line. Furthermore, we observe an inflection point at a genome size of 229 kb, above or below which the percentage of encoded secreted proteins increases with genome size. In contrast, the percentage of encoded intracellular proteins decreases with genome size. We observed no significant correlation between

GC-content and genome size or between coding-density and GC-content for the poxvirus genomes in our data set. Yet, we observed a significant positive correlation between coding density and GC-content for mpox virus isolates (*R*^2^ = 0.799). Further analysis illustrated a rise in both GC-content and coding-density of mpox isolates between 2022 and 2024. Amino acid usage patterns were found to cluster by viral genera, with

Orthopoxvirus, Leporipoxvirus, and Suipoxvirus containing the lowest variation in codon usage frequencies. Tryptophan emerged as the least frequently used amino acid amongst poxviruses. The protein length distribution followed a right-skewed distribution, with the 150-200 amino acid (aa) cohort containing the highest number of predicted proteins. Finally, we observe a significant positive correlation between ITR sizes and duplicate genes (R^2^ = 0.679), highlighting the potential role of ITR expansion in the evolution of poxviruses.

In summary, our findings highlight significant genome architectural relationships that govern the life histories of poxviruses, providing new insights into the constraints and innovations shaping poxvirus biology.

## Materials and Methods

### Genome acquisition and processing

Viral genomes were obtained from NCBI’s GenBank database using the bioinformatics toolkit BIT [10]. Genome metrics, including 1) coding density, 2) genome size, 3) GC-content, and 4) protein sizes, were calculated from each genome’s GFF and genomic FASTA files using custom Python scripts:

### Custom python code

Inverted terminal repeats were predicted using a Python script (https://github.com/Middle-Author-Bioinformatics/PoxVirusCode/blob/main/poxviruses-itr.py), allowing up to 10 mismatches between each terminal repeat sequence. Transcript overlap potential was estimated by comparing the intergenic distance between adjacent open reading frames (ORFs) that are on opposite strands. The code used for this is available: https://github.com/Middle-Author-Bioinformatics/PoxVirusCode/blob/main/dsRNAfinder.py.

### Prediction of gene duplicates

We used ParaHunter to predict gene paralogs and duplicates [14]. ParaHunter performs clustering at the protein sequence level using MMSeq2 to identify clusters of homologous genes. It then uses PAML to identify the degree of divergence within each gene cluster [19]. ParaHunter code is available here: https://github.com/Arkadiy-Garber/ParaHunter.

### Data Analysis

The data was analyzed using a custom Python script generated and modified with ChatGPT. All the scripts are available in the supplemental information.

## Results

### Poxvirus Coding Density

We analyzed a total of 900 complete poxviral genome sequences available on the National Center for Biotechnology Information (NCBI) of classified Poxviridae genera, as listed in the International Committee on Taxonomy of Viruses (ICTV). We observed about a 3.75-fold change in genome size amongst PXV. The largest genome was observed in Carp edema virus (CEVD), measuring 456.8 kilobases (kb), while the smallest belonged to Cetacean poxvirus (CePV), at 121.7 kb [13, 15].

To investigate whether genome size expansion is associated with changes in predicted coding efficiency, we examined the relationship between genome size and coding density, defined as the proportion of the genome occupied by open reading frames. Since poxviruses replicate exclusively in the cytoplasm and do not engage in RNA splicing, there are no known introns. We observed a negative relationship between PXV genome size and coding density (slope = -0.0008, R = -0.466, R^2^ = 0.218) (Figure 1A). The data suggest that larger poxvirus genomes tend to have slightly lower coding densities, possibly due to an increase in intergenic noncoding regions, as overall genome size expansion correlates with the number of predicted protein-coding genes (Figure 2).

**Figure 1:**
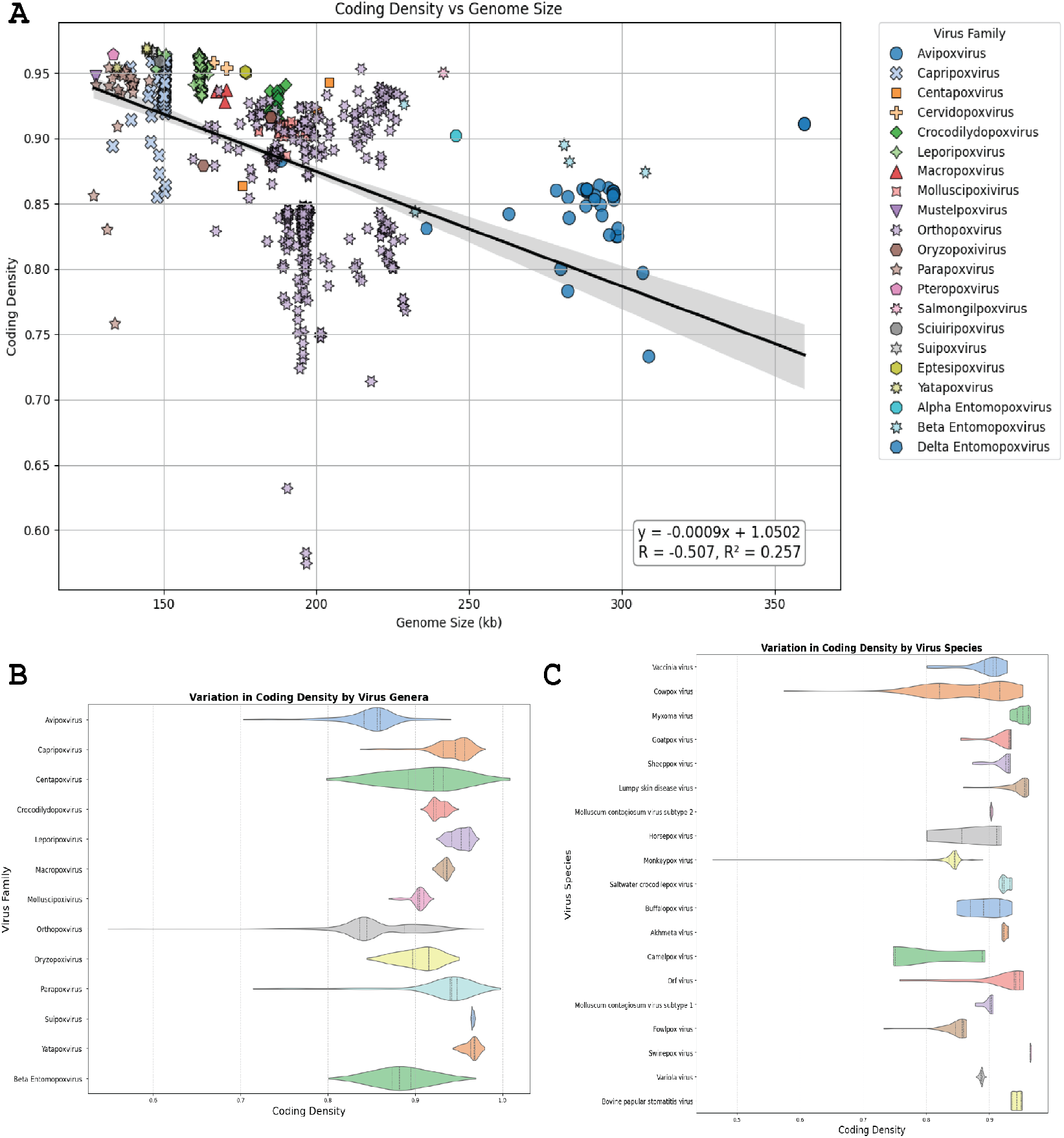
The relationship between poxvirus genome size and coding density. (A) The scatter plot shows coding density as a function of genome size across different PXV species. The y axis displays the coding density (0-1), and the x axis represents genome size in kb. The different colors and shapes represent PXV species. The black line represents the linear regression between coding density and genome size, with the gray-shaded region representing the 95% confidence interval. Violin plot of the variation in virus coding density on the x-axis and virus family (B) or virus species (C) on the y-axis.

**Figure 2:**
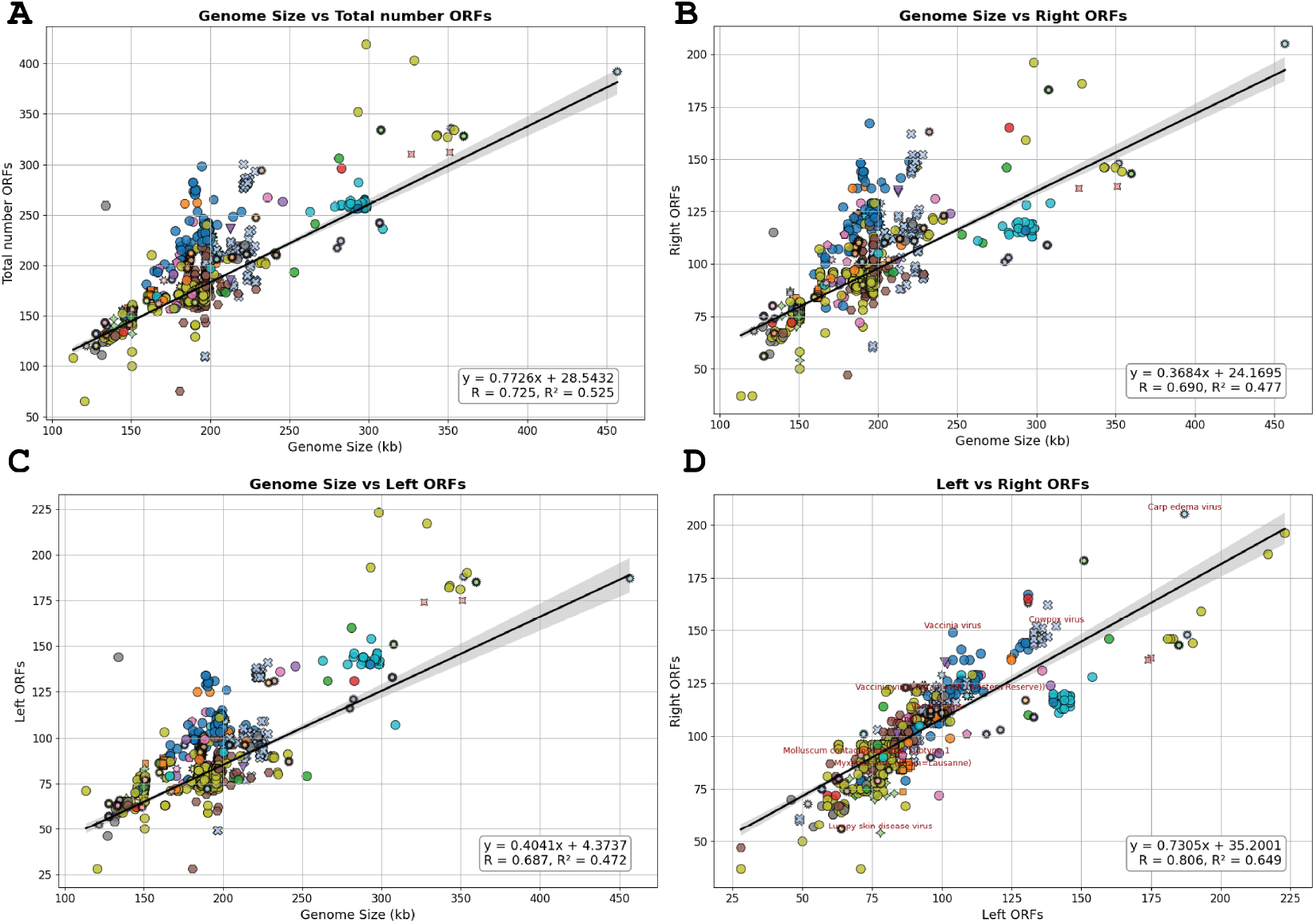
The relationship between poxvirus genome size and number of ORFs. The scatter plot illustrates the relationship between the number of ORFs and genome size across various PXV species. (A) The y-axis displays the number of ORFs, and the x-axis represents genome size in kb. The different colors and shapes represent PXV species. The black line represents the linear regression between coding density and genome size, with the gray-shaded region representing the 95% confidence interval. (B) The y-axis displays the number of right ORFs, (C) left ORFs, and (D) right against left ORF.

Coding densities were conserved across most poxvirus genera, typically ranging from 0.8 to 0.93; however, the genus Orthopoxvirus displayed a broad distribution in coding densities (Figure 1B and 1C). A cowpox virus isolate had the lowest coding density (0.574) in our dataset, whereas a Tanapox virus isolate contained the highest coding density (0.969).

This suggests that the moderate increases in PXV non-coding DNA can be attributed to other biological factors, including immune environments and non-coding elements. The results also illustrate that coding density remains conserved across virus families, although several virus species reveal exceptions, suggesting different life and evolutionary histories within virus species.

### Poxvirus Open Reading Frames

Next, we analyzed the distribution of ORFs across 7,300 poxvirus genomes (genome sizes greater than 100 kb and combined ORFs predicted to encode proteins greater than 50 amino acid). We hypothesized that as the number of PXV genomes increased, the number of ORFs would increase accordingly. Our linear regression analysis supported this prediction, revealing an R^2^ of 0.525, suggesting that more than half the variability in ORF number can be accounted for by genome size (Figure 2). Despite the strong correlation between ORF and genome size, notable strains of Vaccinia virus (VACV), Vaccinia virus Ankara, and Tiantan encoded significantly more ORFs than is predicted, given their genome size. This may reflect strain-specific adaptation and the different biological niches of these viruses.

Given the bilateral coding architecture of the PXV genome, we examined how the left and right ORFs scaled with genome size. We plotted independently the left-ORFs and right-ORFs as a function of genome size, and observed a near-linear relationship with genome size.

To explore whether ORF directionality varies across viral genera, we examined the distribution of ORFs across different viral genera. We observed that specific lineages tend to have higher numbers of either right- or left-oriented ORFs (Figure 2)(Table 3). The data suggest that the distribution of ORFs among PXVs is not symmetrical, indicating lineage-specific preferences.

**Table 1.** List of the top 4 poxviruses with the largest genomes.

| **Subfamily** | **Genus** | **Species** | **Genome Size (Kb)** |
| --- | --- | --- | --- |
| Chordopoxvirinae | Unclassified | Carp edema virus | 456.821 |
| Chordopoxvirinae | Avipoxvirus | Canarypox virus | 359.853 |
| Chordopoxvirinae | Avipoxvirus | Albatrosspox virus | 351.909 |
| Chordopoxvirinae | Avipoxvirus | Shearwaterpox virus | 326.929 |

**Table 2.** List of the top 4 poxviruses with the smallest genomes.

| **Subfamily** | **Genus** | **Species** | **Genome Size (Kb)** |
| --- | --- | --- | --- |
| Chordopoxvirinae | Unclassified | Cetacean poxvirus 1 | 121.769 |
| Chordopoxvirinae | Parapoxvirus | Orf virus | 127.16 |
| Chordopoxvirinae | Mustelpoxvirus | Sea Otterpox virus | 127.876 |
| Chordopoxvirinae | Parapoxvirus | Seal parapoxvirus | 127.941 |

**Table 3.**
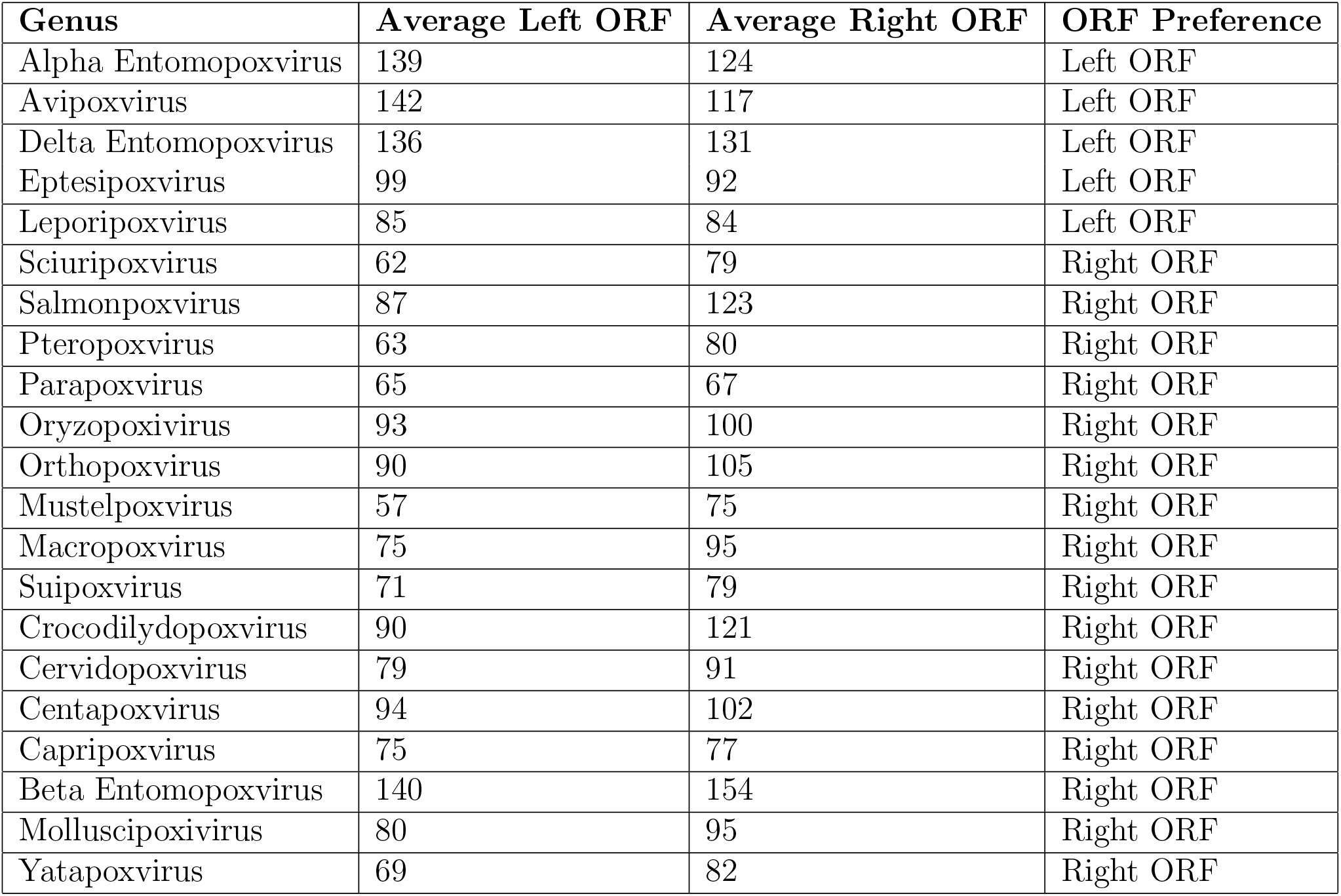
List of the top 4 poxviruses with the smallest genomes.

### Poxviral Transcript Overlap Potential

Virtually all viruses must contend with the issue of virally accumulated nucleic acids and their impact on host immune activation [4**?**]. Poxviruses accumulate double-stranded RNA (dsRNA) during infection, stemming from the complementary base pairing between RNA copies that contain complementary overhangs [5, 1, 2]. Genes located in tandem with each other that have a different orientation in their ORF, for example, genes A and B with orientations A-rORF (right) and B-lORF (left), have the potential to form dsRNA after transcription. The problem arises from polymerase “run-off”–transcribing outside the confines of a gene of interest [5, 1, 2]. Since Poxviruses have both left and right ORFs, and the amounts seem to vary within virus genera. We asked if there is a variation between tandemly located genes on opposite open reading frames (ORFs). Plotting left-right overlap potential spots against genome size, we observed a moderate positive correlation (R = 0.483, R^2^ = 0.234) (Figure 3A).

**Figure 3:**
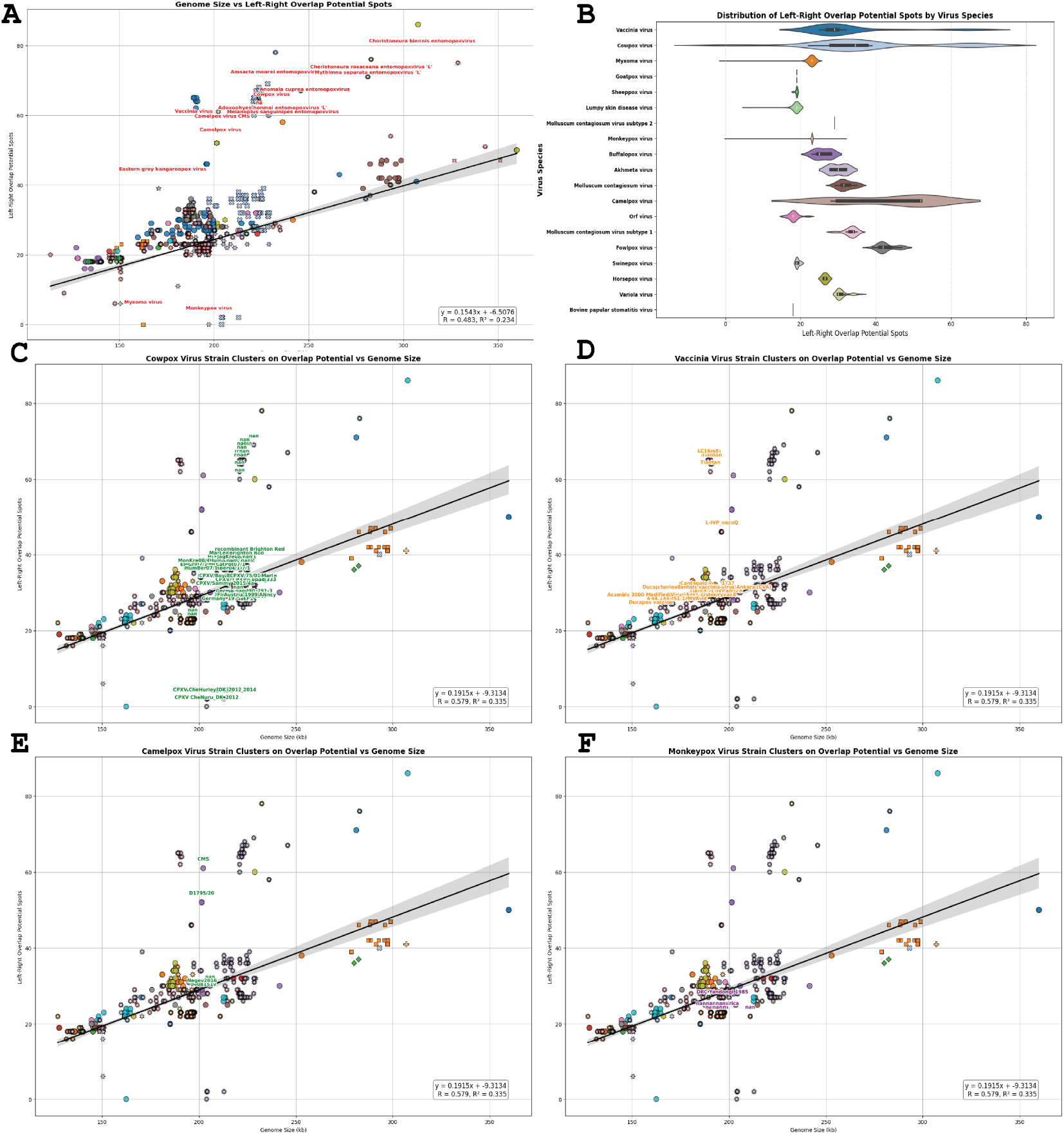
The relationship between poxvirus left-right overlap potential and genome size. (A) The scatter plot shows the number of left-right overlap potentials across different PXV species. The y-axis displays the number left-right overlap potential, and the x-axis represents the genome size. The different colors and shapes represent PXV species. The black line represents the linear regression between coding density and genome size, with the gray-shaded region representing the 95% confidence interval. (B) Violin plot of the variation in virus dsRNA overlap potential on the x-axis and virus family on the y-axis. The scatter plot shows the number of left-right overlap potentials across strains of (C) cowpox virus, (D) vaccinia virus, (E) camelpox virus, and (F) monkeypox virus.

Interestingly, we observed a significant variation around the mean for Orthopoxviruses (Figure 3B). Cowpox viruses displayed more than a 2-fold variation in left-right overlap potential; strains such as EP-4 and Finland 2000-Man exhibited higher overlap potential than the Brighton Red strain. The vaccinia virus overlap potential ranged between 35 and 50 overlap spots, with strains Tiantan and LC16m8 exhibiting the highest overlap potential compared to the Ankara strain. On the other hand, the overlap potential for MYXV was between 32 and 34 bp. The intraspecies divergence in overlap potential, or the lack thereof, might reflect the different evolutionary niches present to poxviruses from various sources.

An unanswered question remains. Since the opposite gene orientation causes immune activation problems, why encode genes opposing ORF in the genome at all? The most logical reason might be that having opposing ORFs maximizes genomic spaces while minimizing intergenic regions. However, there should exist a cost above which further increase in the opposite ORFs negatively affects the fitness of the virus.

### Management of Localization of Encoded Proteins

During virus infection, poxviruses synthesize proteins that localize to both the intracellular and extracellular environments. The extent to which genes that encode intracellular and extracellular proteins vary with genome size amongst poxviruses hasn’t been addressed. Encoded protein sequences were run through Signal-Ip to predict their localization. We plotted extracellular proteins against genome size. The results demonstrated a weak relationship between the extracellular proteins and genome size, with an R^2^ of 0.356 (Figure 4B). Interestingly, we observed a sharp drop in the slope greater than 229 kb. To better resolve the correlation between genome size and protein localization, we fitted a regression line below and above a genome size of 229 kb. For genomes less than 229 kb, we obtained an R^2^ = 0.591, while for genomes greater than 229 kb, the R^2^ was 0.766 (Figure 4C). Next, we analyzed the relationship between intracellular proteins and genome sizes. As predicted, we observed a moderate positive correlation between intracellular protein number and genome size (R^2^ = 0.494), suggesting that larger poxviruses encode fractionally more genes that modulate the intracellular environment (Figure 4A). To get estimates of the fractional investment in intracellular and extracellular proteins, we calculated the percentage of intracellular vs extracellular proteins as a fraction of the total viral proteome. The percentage of extracellular proteins increased nonlinearly with genome size, with a notable inflection point at 229 kb (≤ 229 kb; *R*^2^ = 0.48, *>* 229 kb; *R*^2^ = 0.47). Although we observe a positive correlation for secreted proteins above and below 229 kb, genomes greater than 229kb invested a lower amount of their coding capacity in secreted proteins. It is also worth noting that most poxviruses greater than the inflection point of 229 kb infect non-mammalian species, while viral genomes less than 229 kb are mammal-infecting poxviruses. In contrast, the percentage of cytoplasmic proteins decreased with increasing genome size, both below and above 229 kb (≤ 229 kb; *R*^2^ = 0.48, *>* 229 kb; *R*^2^ = 0.47). This suggests a dilution of intracellular protein investment as PXV genomes grow.

**Figure 4:**
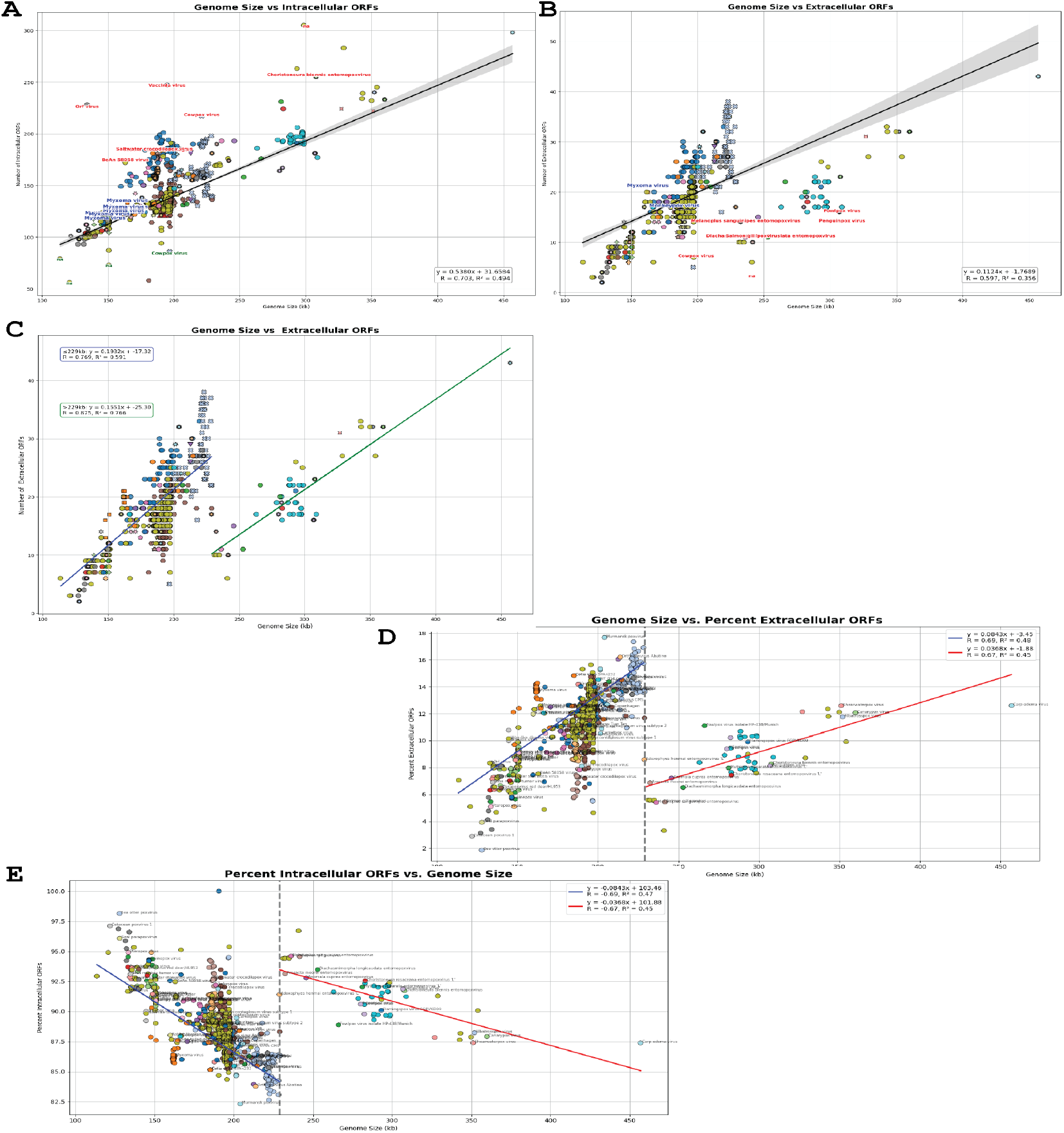
The relationship between poxviral secreted proteins and genome size. (A) The scatter plot shows the number of cytoplasmic proteins across different PXV species. The y-axis displays the number of cytoplasmic proteins, and (B) secreted proteins against genome size on the X-axis. (C) The relationship between poxviral secreted proteins and genome size for genomes below and above 229 kb. (D) Scatter plot of the percent secreted and percent (E) cytoplasmic proteins against genome size. The different colors and shapes represent PXV species. The black/blue/red/green line represents the linear regression between coding density and genome size, with the gray shaded region representing the 95% confidence interval region.

There exists an inverse relationship between intracellular and extracellular protein investment with respect to genome size. This represents a functional shift in the poxviral genome architecture. In summary, larger poxviruses invest a higher percentage of their protein-coding genes in extracellular processes. On the other hand, smaller poxviruses dedicate a correspondingly higher amount of their coding capacity to intracellular pathway. These findings highlight the evolutionary forces that govern the biology of poxviruses, although the molecular mechanisms behind these observations remain to be elucidated.

### Poxviral GC Content and Coding Density

Poxviruses exhibit substantial variability in their GC content, ranging from as low as 25% to as high of 65%. To investigate whether genome size contributes to this variation, we examined the relationship between GC content and genome size across all poxvirus genomes available in our dataset. Our results revealed no significant correlation between the two parameters (R^2^ = 0.046), suggesting that genome size is not an important determinant of GC content variability (Figure 5B).

**Figure 5:**
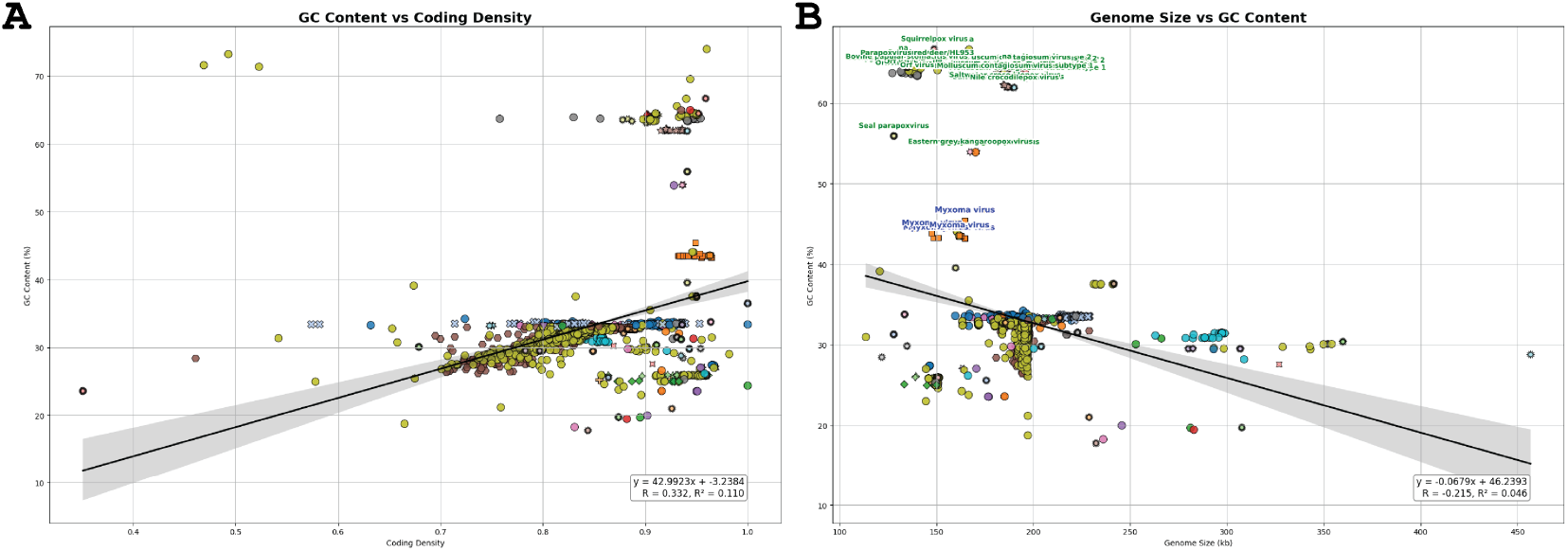
The relationship between coding density and GC content and genome size. The scatter plot shows the GC content across different PXV species and viral coding densities. The y-axis displays the percent GC content against coding density(A) and genome size (B) on the x-axis. The different colors and shapes represent PXV species. The black line represents the linear regression between coding density and genome size, with the gray shaded region representing the 95% confidence interval region.

Given that GC-rich regions are associated with higher protein-coding gene densities in organisms, we examined how GC content affected coding densities. However, similar to our previous observation regarding the correlation between GC content and genome size, we found no correlation between GC content and coding density (R^2^ = 0.110) (Figure 5A). This suggests that GC content cannot account for the variation in coding densities within poxviruses.

Interestingly, we identified that Mpox as a notable outlier in our analysis.

### Mpox GC Content and Coding Density

We further analyzed the observed relation between GC content and coding density for the mpox virus. Mpox virus genomes exhibit a robust positive relationship between GC content and coding density, with an R^2^ of 0.799 (Figure 6A). This suggests that most of the variation in GC content can be attributed to coding density among mpox samples. Since most of the mpox samples in our data were deposited during pandemic-related events, we further separated the GC content and coding density relationship by the year in which the sequence was deposited. Samples deposited between 2009 and 2020 were not plotted because they contained a small number of deposited genomes. A total of 3694 mpox virus genomes deposited between 2022 and 2024 were analyzed to determine how the GC content relates to coding density. Linear regression analysis performed for each year reveals a gradual increase in coding density and GC content for the samples deposited between 2022 and 2024. Genomes with higher GC content invariably contained higher proportions of protein-coding genes. The magnitude of the relationship weakened over time, R^2^ = 0.84 in 2022 to R^2^ = 0.60 in 2024. This attenuation coincides with a tighter GC content and reduced variance in coding densities in 2024, suggesting an increase in sample homogeneity.

**Figure 6:**
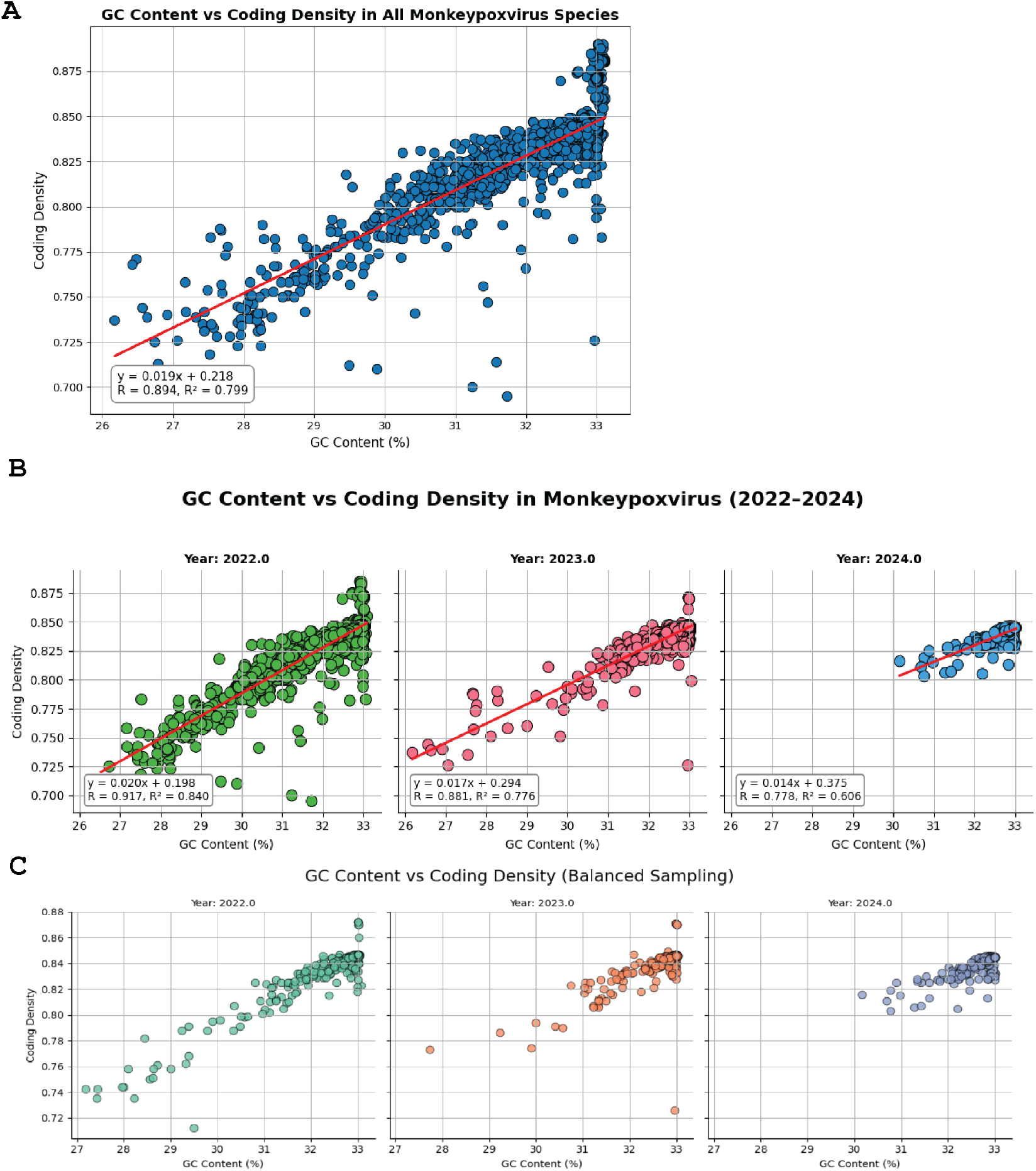
The relationship between coding density and GC content. The scatter plot illustrates the relationship between coding density and GC content. (A) The y-axis displays the percent coding density, and the GC content is represented on the X-axis. (B) Change in coding density and GC content by year. The color represents Mpox genomes. The black line represents the linear regression between coding density and genome size, with the gray-shaded region representing the 95% confidence interval.

Our results for mpox virus samples sequenced between 2022 and 2024 reveal a positive correlation between GC content and coding density, with a gradual increase in coding density observed from 2022 onwards (Figure 6B).

### Poxviral Codon Usage

We analyzed the amino acid usage bias across 7,323 poxvirus species, spanning all poxvirus genera. To reduce dimensions, the data was examined using a principal component analysis plot (PCA), and the relative frequencies of the standard 20 amino acids were utilized to create the PCA. The analysis identified 2 dominant axes accounting for most of the variation. PC1 accounted for 55.7% of the variation, with hydrophobic –Isoleucine (I), Leucine(L), Valine (V)–and polar/charged–Cysteine(C) and Tryptophan(W)–residues mainly contributing to the variations. In contrast, the variation in PC2 (18%) was accounted for by charged residues: Lysine (K), Arginine (R), Glutamate (E), and Aspartate (D). The plot illustrates that Molluscipoxvirus contains the most significant amounts of hydrophobic residues, while Salmonpoxvirus contains the highest abundance of acidic residues. To examine the genus-wide diversity in codon usage, we compared PC1 and PC2 with those of poxvirus genera. We found that Sciuropoxvirus, Yatapoxvirus, Betaentamopoxvirus, and Centapoxvirus exhibited the highest variation in codon usage bias. On the other hand, Orthopoxviruses, Leporipoxviruses, and Suipoxvirus showed the lowest variation in codon usage (Figure 7).

**Figure 7:**
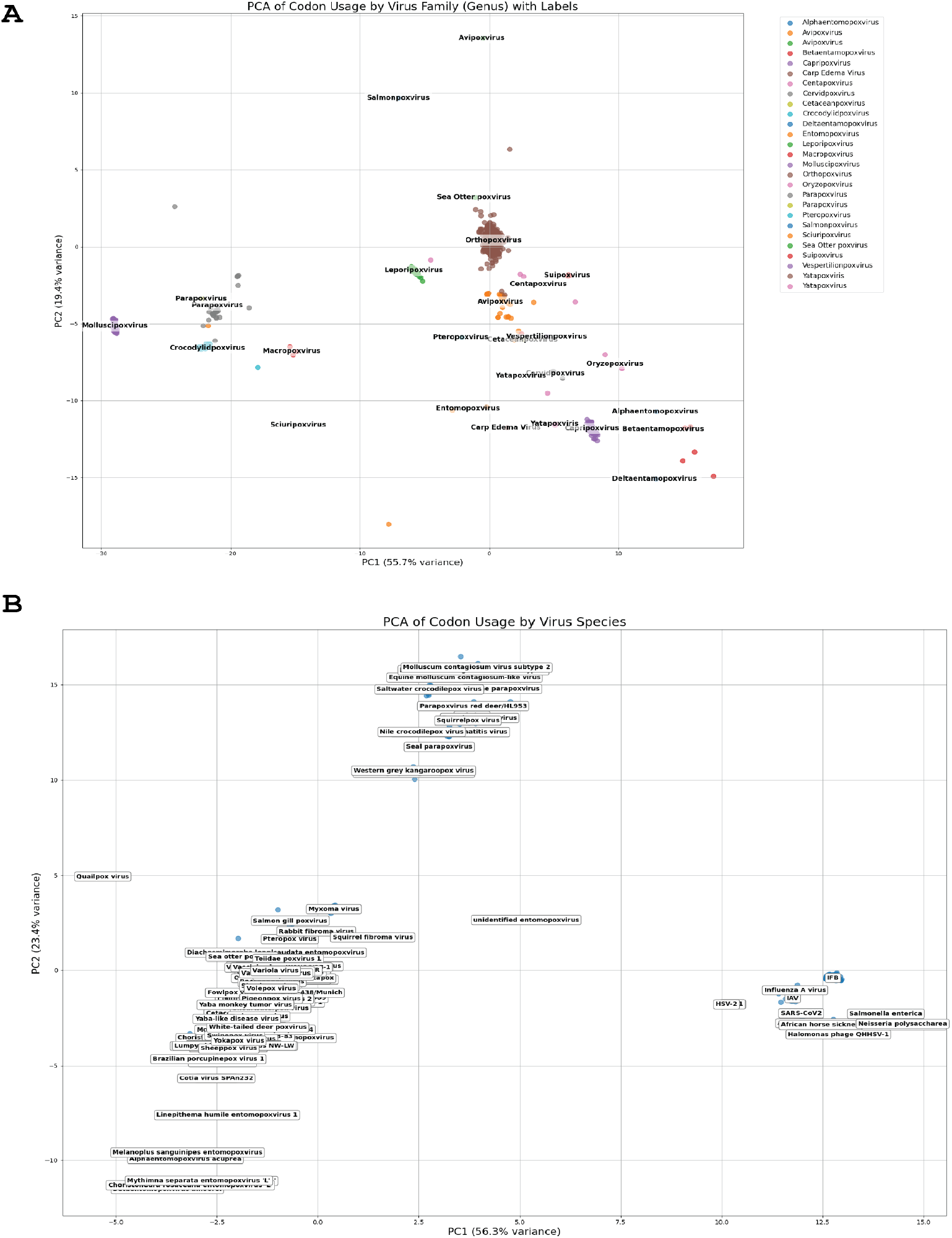
PCA of codon usage bias amongst Poxviridae. (A) The different points represent the codon usage across unique poxvirus genera, represented in different colors, and (B) Non poxvirus species included in the PCA plot

We next examined the mean amino acid usage frequency on a heat map and observed that L and I are the most abundant residues among poxviruses (mean frequency of 11–15%). At the same time, W and C are the rarest residues (Figure 8).

**Figure 8:**
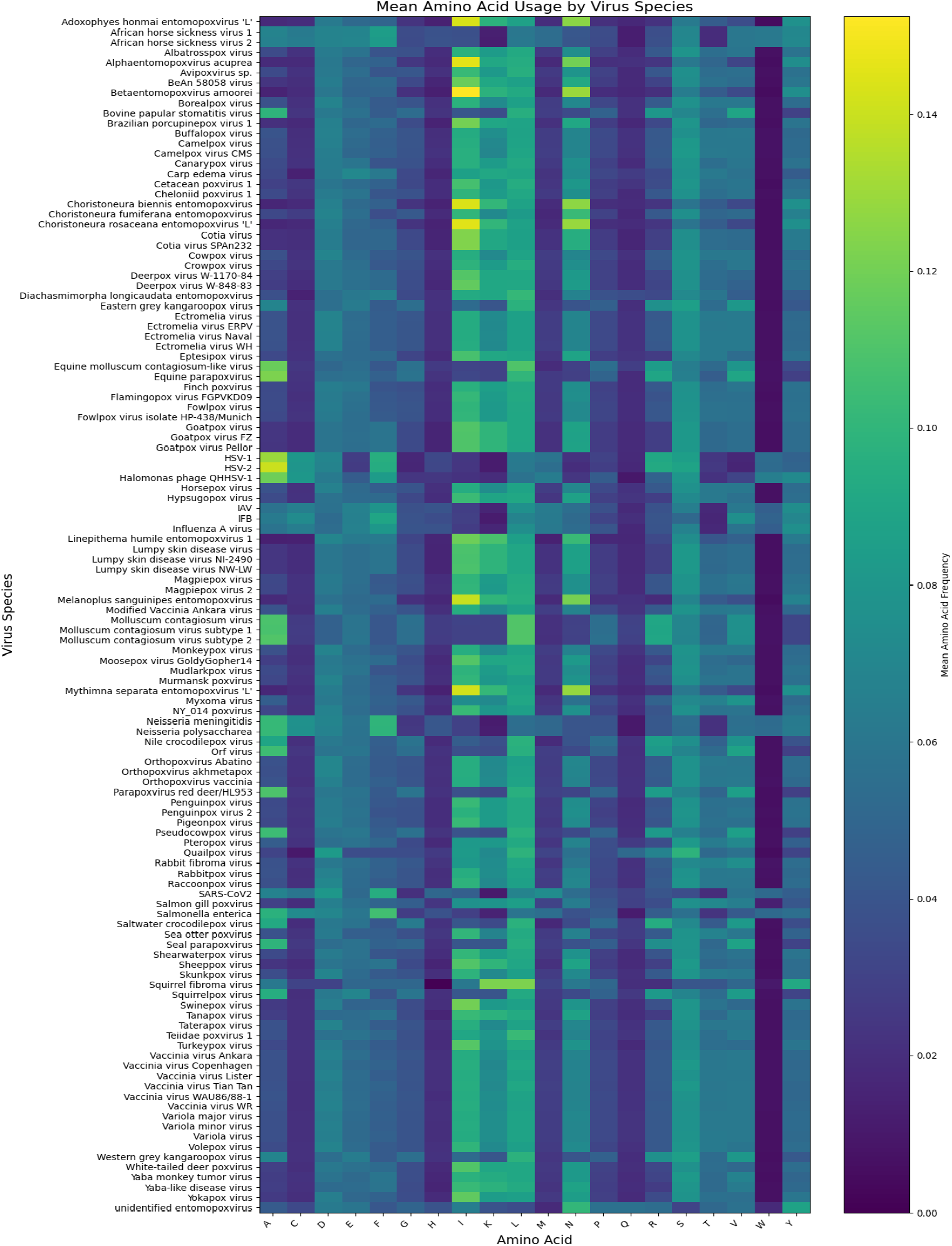
Heatmap of amino acid usage amongst Poxviridae. The column displays the standard 20 amino acids at varying intensities, with yellow representing the highest amino acid frequencies and purple representing the lowest. The rows stand for the poxvirus species.

The observed genus-specific codon usage variation in poxviruses suggests diverse evolutionary strategies with preferences and avoidance of specific amino acids.

### Poxviral Protein Size

To access the distribution in protein sizes amongst poxviruses, we examined analyzed about 1.3 million proteins from the poxvirus species present in our dataset and observed a right-skewed global distribution of protein length (median = 234 aa, mean = 306 aa) Most species show a single peak around 150 – 250 aa with a long low density tail extending beyond 1000 aa (Figure 9A). The largest protein in our data set belonged to a surface glycoprotein of Carp edema virus (2772 aa). Next, we ranked species by deviation from the global distribution. The 10 most deviant species are highlighted in distinct colors (Figure 9B). The results suggest that a majority of poxviral proteins are less than 500 amino acids (aa), underscoring the need for smaller and more compact proteins.

**Figure 9:**
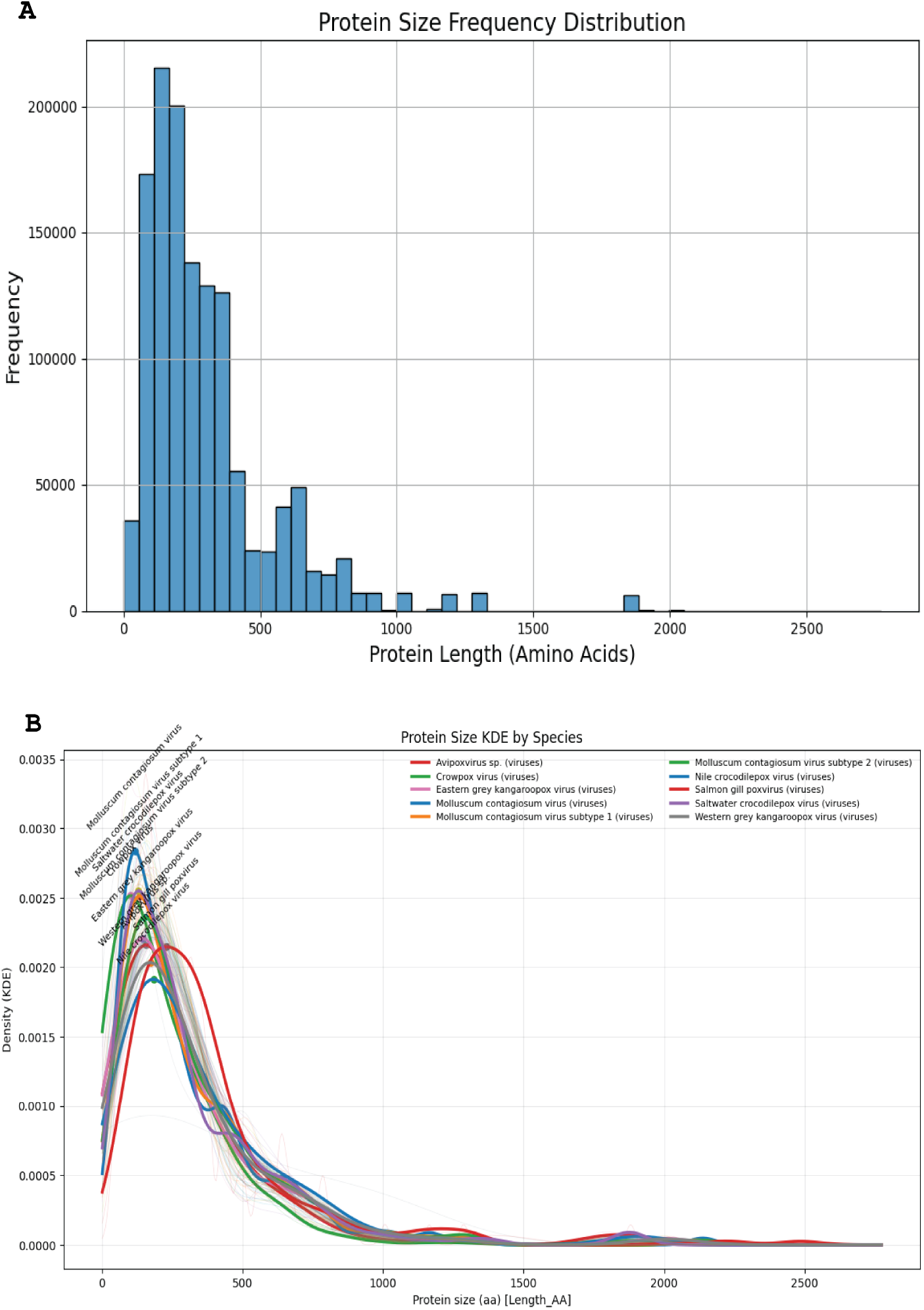
Protein Frequency Distribution Amongst Poxviruses. (A) Frequency distribution of amino acid sequence size for different Poxvirus proteins. (B) Kernel density estimates of poxviral protein lengths. The gray line shows all species. Colored lines indicate the top 10 most deviant species based on standard deviation and distance from the global distribution.

### Poxvirus ITR Size and Duplicate Gene

To investigate the genomic architectural features that influence ITR size, we compared published ITR sizes from annotated poxviruses with computationally derived ITR sizes to verify the agreement between the two values. The regression line of derived versus empirical ITR size yielded an R^2^ value of 0.83 (Figure 10A). Across the 106 derived samples in our dataset, 55 samples had empirical values that showed a weak positive correlation with genome size (R^2^ = 0.139) (Figure 10C). On the other hand, the derived ITR size showed no correlation with virus genome size (Figure 10B).

**Figure 10:**
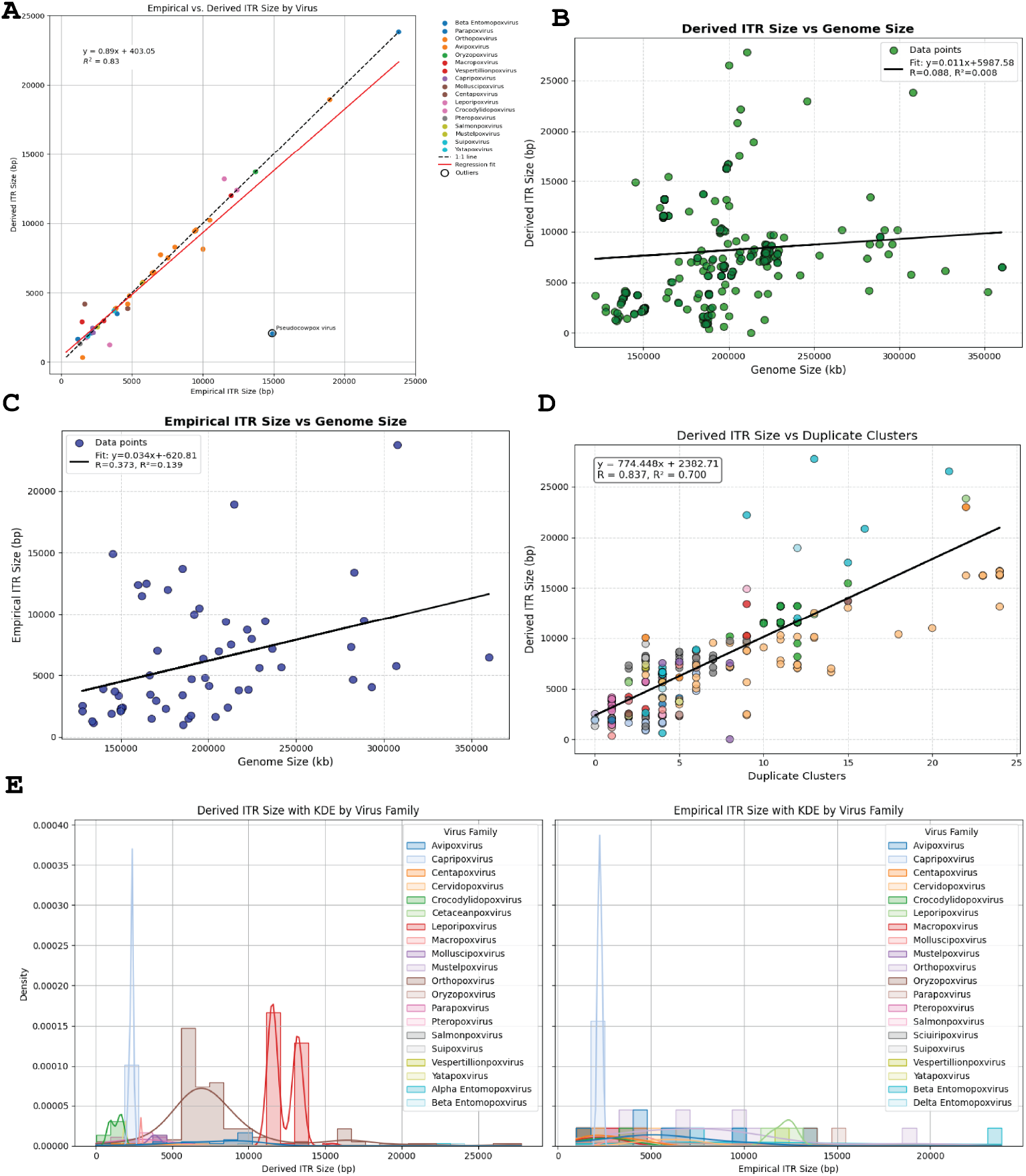
The relationship between empirical, derived ITR size, duplicate genes, and genome size. (A) Scatter plot of the empirically derived ITR vs those calculated using a computational pipeline (derived ITR). Each plot represents a genome colored by virus genera. The orange line represents the regression line. (B) Scatter plot of computationally derived ITR sizes against genome length. The solid black line represents the regression line. (C) Scatter plot of empirical ITR sizes against genome length. The solid black line represents the regression line. (D) Derived ITR size against the duplicate gene cluster. Scatter plot of derived ITR sizes against duplicate genes. Each plot represents a derived ITR size and number of duplicate genes for a virus species. The solid black line represents the regression line. (E) Frequency distribution of derived and empirical ITR sizes across viral genera. Kernel density estimates of poxvirus ITR sizes grouped by virus genera for both derived and empirical viral ITR sizes.

One proposed mechanism by which poxviruses adapt is via the genomic accordion. During the inflationary phase of the accordion, poxviruses amplify maladaptive viral inhibitors to counteract the host immune response, while sampling the mutational space for a more optimal inhibitor. The subsequent gain of an adaptive viral inhibitor coincides with genome contraction to offset the cost of a larger genome [6, 7]. We hypothesized that duplicate genes might correlate with a signature of adaptation, and ITRs are known to be involved in poxvirus adaptation. To further investigate this, we plotted ITR sizes against duplicate gene clusters (Figure 10D) and observed a moderate positive correlation between these two variables (R^2^ = 0.679), suggesting that ITR expansion may be driven at least in part by gene duplication events. Lastly, we observed no specific patterns in the distribution of poxviral ITR sizes, but rather a family-specific clustering. Our data suggest that viral adaptation might drive the expansion of ITR sizes.

## Discussion

Poxviruses are large dsDNA viruses that vary widely in genomic size, yet there are no studies quantifying their genomic architecture. In this study, we identify coordinated relationships among genome size, coding density, ORF quality and quantity, protein localization, GC content, codon usage, protein size spectra, and ITR architecture by analyzing hundreds of genomes. Although each trend is modest in isolation, together they provide an outline of how poxviruses grow their genomes, primarily by the addition of accessory proteins.

We observed a modest decline in coding capacity as genome size increased (R^2^ = 0.25). A reason for this might be the increase in non-coding intergenic DNA as viral genomes inflate in size. Alternatively, the non-coding DNA may provide a structural or regulatory role during infection, as an adaptation. This moderate increase in non-coding DNA raised a question about the cost of non-coding material in the poxvirus genome. To what extent can natural selection regulate the accumulation of non-coding DNA?

A notable trend in our results is the correlation between the potential for transcription overlap and genome size. Because poxvirus transcription termination can be leaky, expanded genomes may face an obstacle in the form of increased transcript overlap potential and the corresponding innate immune activation. Although empirical work is needed to validate the importance of the genome architecture in driving the accumulation of viral transcripts, our data highlight the relevance of the viral genome structure in dictating viral transcripts.

Another salient point in our data is the change in slope for protein-localization fraction at 229 kb. Below this point, the percent secreted proteins increases with genome size, and the cytoplasmic fraction correspondingly declines. Above 229 kb, the relationship persists but with a different slope. We interpret this as evidence that as genomes grow into the 230 kb range, lineages invest more in extracellular accessory proteins while intracellular proteins saturate. Although empirical evidence is lacking, this may account for the broader tropism within viral genera, such as the Orthopoxviruses.

In mpox, we observed a strong correlation between GC content and coding density. Both GC content and coding density rose between 2022 and 2024. Even after controlling for equal sample numbers, the rise in GC content and coding density for mpox remained significant, suggesting that short-term shifts in GC content are unlikely to be driven by de novo mutation or sequencing errors. A plausible explanation for the trend might be clade-specific deposition or viral host adaptation.

Across 1.3 million proteins, we observed a right-skewed protein distribution with a strong mode at 150–200 amino acids. This distribution suggests that poxviral proteins are small modular proteins. The codon usage clustered by virus family is consistent with viral phylogenetic and host adaptation histories. Tryptophan and cysteine are relatively rare, possibly due to the metabolic and immune-related costs associated with these amino acids.

We observe a strong correlation between ITR size and duplicate gene clusters, supporting the notion that poxviral ITRs serve as centers for viral innovation. The virus-specific distribution of ITR sizes suggests that lineage-specific pressure drives the growth and distribution of ITRs.

Several observations are consistent with the idea that genome expansion is not random but rather enriches advantageous functions, such as secretory immune-modulatory proteins. The resulting shift in protein localization,

ITR size, protein size spectra, and codon usage is coherent across virus families, yet they maintain lineage-specific signatures. These patterns provide a quantitative framework for interpreting poxvirus evolution and a playbook for engineering poxviral vectors with predictable behaviors.

## Acknowledgments

We want to thank Michael Lynch for suggestions regarding the manuscript.

